# Limb regeneration is blastema dependent in a ladybird beetle, *Cheilomenes sexmaculata* (Fabricius)

**DOI:** 10.1101/2023.08.28.555044

**Authors:** Saumya Rawat, Shriza Rai, Geetanjali Mishra

## Abstract

Holometabolous insects undergoes metamorphosis which involves an intercalated pupal stage between larva and adult. The body plan of the adult is established during pupal stage and larval systems are de-differentiated and reorganized in insects undergoing complete metamorphosis. In ladybird beetles, limbs amputated in larval stages are regenerated in adults. This occurs during pupation. Given that changes in pupa are akin to embryogenesis, does the lost limbs are redeveloped as a part of metamorphosis or has some pre-patterning initiated prior pupation? To study this we hypothesised that limb regeneration in a holometabolous ladybird beetles, *Cheilomenes sexmaculata* is the result of the recapitulation of the embryonic gene programs in the pupa. To test this, we exposed third larval stages of *C. sexmaculata* to amputation and prior to amputation the tissues on the amputated site was scraped off every 24hrs. It was found that limb regeneration does not occur in the treatment where scrapping was done. Assuming that these epidermal cells correspond to blastema, for limb regeneration in ladybird beetles blastema is essential and does not occur in its absence.

## Introduction

More than 80% of insects undergo complete metamorphosis or holometaboloy with an intercalated pupal stage between larva and adult (Engel and Grimaldi, 2005). The pupal stage, which is largely immobile, involves extensive remodelling of organs and tissues, resulting in rebuilding of entire body plan (Rolff et al., 2019). Many metabolic and developmental genes have been reported to revert to embryonic-like state during metamorphosis. Ozerova and Gelfand (2022), hypothesised that there is recapitulation of the embryonic expression program during metamorphosis and demonstrated an increased similarity of pupa to embryonic stage via metanalyses of numerous datasets of holometabolous insects.

Ladybird beetles are holometabolous insects that undergoes limb regeneration (Wang et al. 2015, Saxena et al. 2016, Michaud et al., 2020). An amputated limb is regenerated when larva metamorphosizes into adult. The capacity of limb regeneration has been reported to be broadly conserved in coccinellids with 15 out of 16 species of ladybird beetles exhibiting limb regeneration with delayed pupal duration. Regeneration neither occurred from one moult to another nor the amputated legs in adults showed regeneration (Wang et al., 2015; Saxena et al., 2016) and was evident only during pupal development. An activation of genes responsible for regeneration probably occurred in pupal stage (Michaud et al., 2020; Zhong et al., 2023). Possibly the cells initiate differentiation during pupation forming imaginal primordia, contributing to the development of adult structures and maintain pluripotency similar to stem cells (Truman and Riddiford, 2002; McClure and Schubiger, 2007). In *Drosophila*, regeneration occurs via imaginal disc formation while in *Gryllus* regeneration involves blastema formation; however, the signalling involved during regeneration is similar in both (Das. 2015). Having said that it is yet to be explored if redevelopment of lost limbs in pupa is a part of metamorphosis or involves the process of regeneration via blastema formation. Since pupal stage involves whole-body reorganization; most larval tissues and organs are respecified including epidermis (Riddiford 1978) and muscles (Lubischer et al 1999), it was hypothesized that limb regeneration in holometabolous insects like ladybird beetles, is the embryonic recapitulation of the gene programs in the pupa and is therefore independent of blastema. Therefore, this experiment will help us to understand if the ability to form a blastema is directly related to the ability to regenerate lost legs or is a part of metamorphosis.

Ladybird beetles are known globally for their biological potential. During mass rearing, it is not uncommon for larvae to incur nonlethal injuries including appendage loss owing to fierce cannibalism exhibited by these beetles (Michaud 2003). Current study will be helpful in increasing our understanding of limb regeneration in non-model insects like ladybirds and offer better insight into the evolution of regeneration.

## Materials and Methods

### Study Organism

*Cheilomenes sexmaculata* (Fabricius) was selected for this study because it exhibits limb regeneration (Saxena et al., 2016) and is widely distributed in the local agricultural fields of India.

### Stock Maintenance

Adults of *C. sexmaculata* (n=20) were collected from the fields of Lucknow, India (26°50’N, 80°54’E), and were reared for two generations in the laboratory. They were provided with an *ad libitum* supply of *Aphis nerii* Boyer de Fonscolombe (Hemiptera: Aphididae) on the host plant *Calotropis procera* (Family: Apocynaceae) commonly known as milkweed aphid which is found locally in abandoned lands. Stock culture of beetles was done in a 1 litre plastic beaker and kept in temperature-controlled room set to 27±2°C, 65±5% R.H. Mated females were removed from beakers and transferred to plastic Petri dishes (9.0×2.0 cm) and provisioned *ad libitum* aphids. The eggs laid were collected every 24h, and observed for hatching and the first instar larvae were used in the experiment.

### Experimental Design

120 newly hatched larvae from the stock were placed in two sets of 40 each and kept singly in Petri dishes (9.0×2.0 cm). In the first set, amputation was performed in third larval instar (L3) stage. At first, the larvae were chilled individually for 5 min in a refrigerator and were amputated at the proximal end of the femur of the right foreleg. Amputation was performed under a stereomicroscope (Magnus)with the magnification 16X with the help of a micro-scalpel. After 24hrs of amputation, the epidermal growth at the proximal position of amputation site was scraped off using a micro-scalpel under the stereomicroscope (Magnus). This was repeated after every 24hrs till the larva underwent pupation. In the second set there was amputation in L3 stage but no scraping off was performed while in third set there was no amputation and was called the control. All three sets were maintained on *ad libitum* supply of *A. nerii*, until the emergence of the adult. Petri dishes were cleaned every 24 hours throughout the experiment to maintain the prey quality and to avoid any microbial contamination. Larvae were examined daily to record their larval stage developmental durations, pupal duration and post amputation developmental duration, for amputated (scraped), amputated (unscraped) and control beetles.

### Statistical Analysis

Data sets were first tested for normality using the Kolmogorov-Smirnov test and were found to be non-normal for larval stage developmental durations, pupal duration, and post amputation developmental duration of Control, Scraped and Unscraped beetles as treatments. On finding the data non-normal, data were subjected to Generalized Linear Model (GLM) with Gamma log link. Treatments were taken as fixed factors and larval stage developmental durations, pupal duration, and post amputation developmental duration were taken as dependent factors. This analysis was followed by a comparison of means using Wald Chi-Square test. The analysis was done using IBM SPSS version 20 statistical software.

## Results and Discussion

Out of a total of 40 replicates, only 6 regenerated the amputated leg, while the rest did not regenerate (Figure 1). The everyday scraping off of the tip of the regenerating leg corresponds to removal of blastema. Thus, our results demonstrated the importance of blastema for the regeneration of lost limbs in ladybird beetles. The 6 individuals that showed some level of regeneration of the leg amputated in the larval stage, could be the failure to completely scrape off the blastema. Thus, even some remaining cells of blastema initiated regeneration but were not enough to lead to complete development of the lost leg.

**Figure 1:**
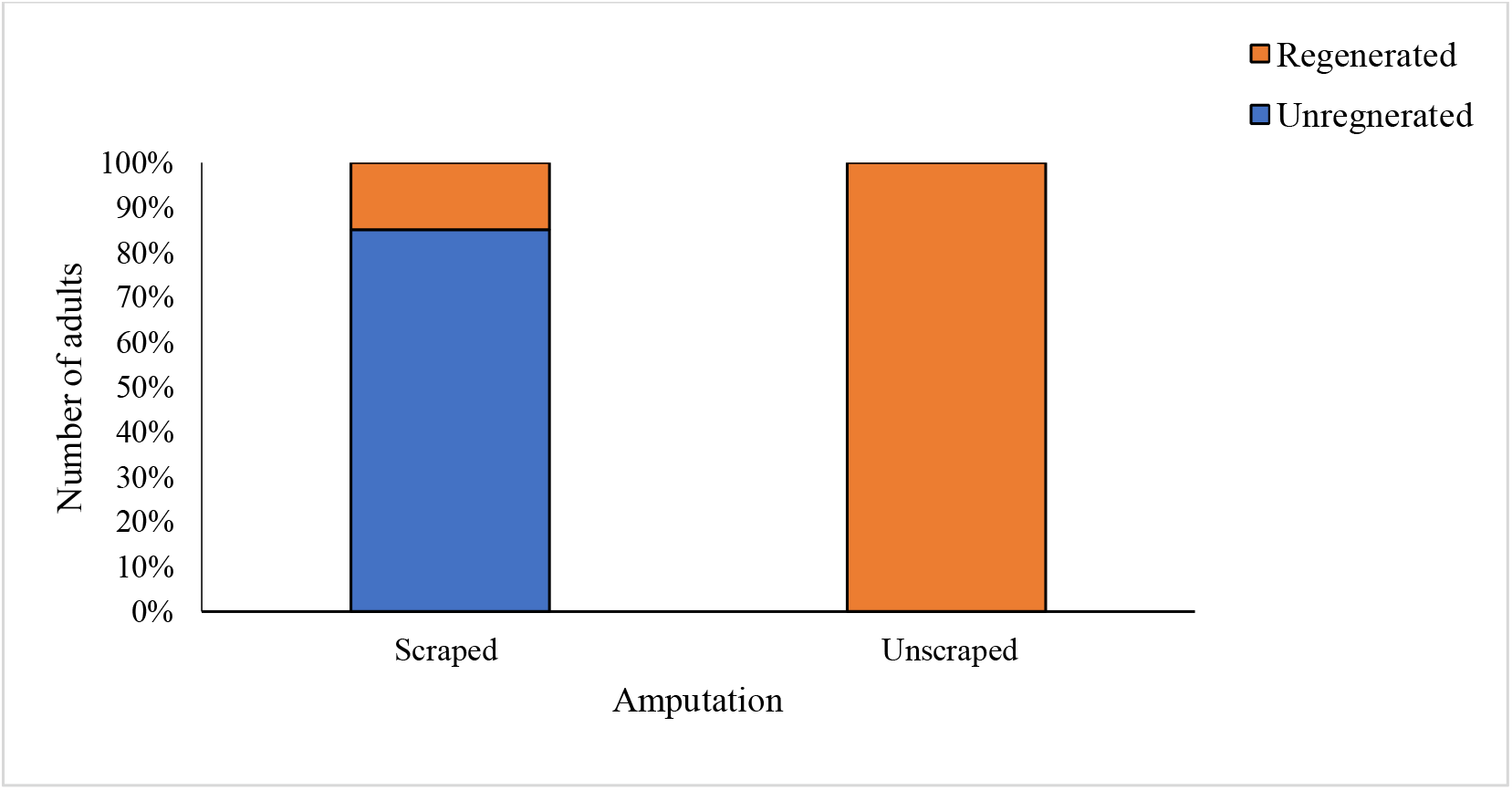
Effect of scraping off of proximal portion of amputation site on regeneration in comparison to the unscraped larvae of *Cheilomenes sexmaculata*.

The proximal part of the amputated site, gets covered by a scab of epidermal cells. The removal of this scab probably resulted in the removal of blastema because of which there was no regeneration in the treatment that involved scraping. This demonstrates the blastema dependency of limb regeneration in *C. sexmaculata*. Das (2015) reported that limb regeneration in insects is initiated via scab formation, followed by migration of epidermal cells at the site of amputation and these epidermal cells known as blastema, acted as the precursors of the regenerating cells. Various cellular processes and signalling unfold in blastema ultimately specifying the cells to give rise to the formation of lost structures (Zhong et al., 2023). There is a combination of genetic factors forming a ‘prepattern’, which presages the final differentiated structure. For example, several genes like *dachshund* (Dac) aid in regeneration and their absence resulted in abnormally shortened legs (Lee et al., 2013). Dac silenced in amputated larval leg, affected the patterning of the adult leg, indicating that larval appendage morphology is inherited by the adult appendages.

Amputation of foreleg led to delayed post amputation development duration with maximum delay in unscraped treatment (Table 1). Pupal duration was found to be maximum in unscraped treatment followed by scraped treatment and control. Interestingly, the pupal duration in scraped treatment was similar to control. This difference from previous studies where the maximum delay of amputated beetles was reported in the pupal stage (Wang et al., 2015; Saxena et al., 2016; Michaud et al., 2020; Rai et al., 2023), can be attributed to the absence of blastemal cells for the initiation of regeneration and hence no regeneration occurred in pupa, therefore no delay. While delay in the developmental duration of the fourth larval stage of the amputated treatment demonstrated the attempt to form the blastema via cellular proliferation, with no blastema in the larvae entering pupation there was no initiation of regeneration. Since, regeneration in ladybird beetles occurs during pupal duration (Wang et al., 2015; Saxena et al., 2016; Michaud et al., 2020; Zhou et al., 2021), with no regeneration taking place in the pupa, the amputated larvae of scraped off treatment showed no delay in the pupa. This explains that the delay in the pupa is indicative of regenerative processing occurring inside the pupa.

**Table 1:**
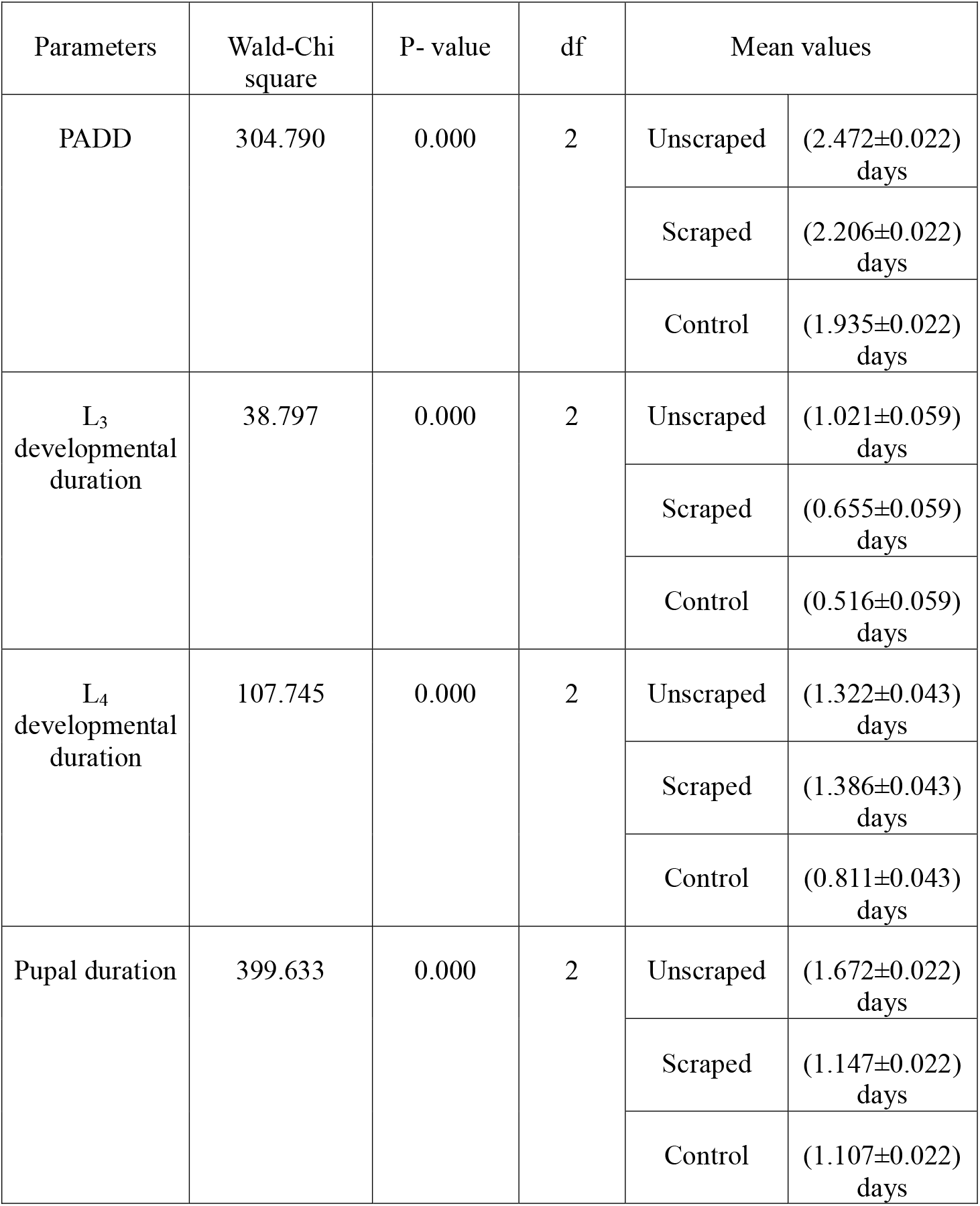
Results of GLM with Gamma log link depicting the effect of amputation on Post amputation development duration (PADD), larval development durations (L_3_ and L_4_) and pupal duration in *Cheilomenes sexmaculata*, along with their mean values.

Our hypothesis that the legs amputated in the larval stages, are redeveloped during metamorphosis as a part of embryonic recapitulation of the gene programs in the pupa, was rejected. This study concluded that (i) scraping of scab formed at the proximal position of the amputation site, prevented the formation of blastema, (ii) the ability to form a blastema is directly related to the insects’ ability to regenerate lost legs, and (iii) the delay in pupa is indicative of undergoing regeneration processes. Thus, limb regeneration in ladybird beetle *C. sexmaculata* is blastema dependent and the lost structure is reconstructed from blastema. Further studies are needed to understand the role of biotic and abiotic factors on blastema development. Transcriptome analysis of pupa with blastema and without blastema is needed to further understand the molecular aspect of limb regeneration in ladybird beetles.

## Acknowledgements

GM acknowledges SERB DST (F.No. CRG/2020/001095 via letter no. BP-2020-20-3308 dated December 30, 2020. SR acknowledges UGC Fellowship by University Grants Commission, New Delhi, India (F.No.16-9(June 2019)/ 2019(NET/CSIR)) dated August 09, 2019.

